# Evaluating the effects of archaic protein-altering variants in living human adults

**DOI:** 10.1101/2024.07.05.602242

**Authors:** Barbara Molz, Mikel Lana Alberro, Else Eising, Dick Schijven, Gökberk Alagöz, Clyde Francks, Simon E. Fisher

## Abstract

Advances in paleo-genetics allowed the identification of protein-coding changes arising on the lineage leading to *Homo sapiens*, by comparing genomes of present-day and archaic hominins. Experimental validation of the potential impact of such changes has so far been restricted to functional assays and model organisms. Large-scale biobanking now makes it possible to identify present-day carriers of archaic alleles and to directly assess phenotypic consequences in living adults. We queried exomes of half a million people in the UK Biobank at 37 genomic positions with supposedly fixed human-specific protein-coding changes. This yielded 103 carriers at 17 positions, with variable allele counts across ancestries. Contrasting carriers and non-carriers of an exemplary archaic allele in *SSH2*, we observed no deviation from the norm in a range of health, psychological, and cognitive traits. We also identified 62 archaic-allele carriers for a *TKTL1* missense change, previously shown to have large effects on cortical neurogenesis in brain organoids and animal models. Carriers did not show differences in relevant anatomical brain measures, and a substantial proportion had college/university degrees. This work offers an empirical demonstration of how large-scale biobank investigations of living adults can transform our understanding of human evolution. The findings challenge the notion of fixed human-specific genomic changes, highlight that individual interrogation of relevant sites is unlikely to yield major insights into the emergence of complex human traits, and emphasise the importance of including diverse ancestries when investigating origins of our species.

## Main Text

Understanding the origins of modern humans and how our ancestors developed sophisticated cultural, social and behavioural skills has been a central issue for many fields of science (*1*–*3*). While latest research is gradually reaching a consensus that cognitive capacities of Neandertals were greater than previously appreciated, the question remains why *Homo sapiens* outlived its archaic cousins and was able to migrate all across the globe (*2, 4*–*6*). Advances in high-throughput DNA sequencing and the availability of three high quality Neandertal genomes (*7*–*9*) enabled comparative genomic approaches, opening up new ways to reconstruct aspects of the evolutionary history of *Homo sapiens*. In particular, such approaches yielded catalogues of missense variants (changes that substitute one amino acid for another in an encoded protein) that occurred after *Homo sapiens* split from its common ancestor with Neandertals ∼600,000 years ago, and that reached (near) fixation on our lineage. These human-specific fixed derived alleles have been hailed as promising entry points for explaining human origins, given their enrichment in genes that are relevant for human-specific traits and involved in cortical development and neurogenesis^1,2,7^.

Since a missense variant can potentially arise and spread through a population without any consequence to properties or functions of the encoded protein, experimental validation is crucial to determine the functional significance of derived alleles. In a prominent example, Pinson et al. (*10*) investigated the impacts of a lysine-to-arginine substitution in human *TKTL1* (chrX:154,315,258; G->A) by comparing the archaic and derived alleles using genome-edited cerebral organoid and *in vivo* models, as well as in primary brain tissue. The authors observed substantial differences between samples carrying the Neandertal and *Homo sapiens* versions of *TKTL1* in basal radial glia abundance and neurogenesis, and suggested that the modern human-derived allele might have played a key role in evolutionary expansion of the brain’s frontal lobe. However, despite the multiple strengths of cerebral organoids for modelling events in early embryogenesis (*11*), cellular diversity and transcriptomic programmes of these models do not fully recapture human brain development, and lack insights from diversity across genetic ancestries (*12*). Similarly, expression of “humanised” genes in primary brain tissue of non-human species may lead to non-specific artefacts (*12*–*14*), due to inter-species differences in genetic background. Thus, the actual consequences of any such modern human-derived genetic changes may be more complex than those which can be observed in cellular/animal models (*15*).

A complementary approach for evaluating broader biological impact that has only recently become feasible, depends on the identification of present-day living carriers of archaic alleles at genomic positions that differ between modern humans and Neandertals (*3*). Indeed, databases like gnomAD highlight the existence of individuals carrying these archaic single nucleotide variants (aSNVs) (*12*), albeit in low numbers. With availability of large-scale biobanks with exome sequencing and trait data it is now possible not only to detect aSNVs in living humans, but also to investigate putative phenotypic consequences in a way that could not be done before.

In this study, we used the UK Biobank (UKB), a large-scale population resource with both exome and dense phenotype data available from around half a million individuals (*16*). This offers a unique opportunity to i) determine the frequencies of present-day aSNV carriers, and ii) assess how phenotypic profiles of carriers of the archaic allele compare to individuals that are homozygous for the derived present-day human allele. We focused our efforts on a catalogue of putative fixed genomic positions established from a prior survey of potential human-specific changes (*2*) and searched for carriers of ancestral alleles among UKB participants. To gain insight into the phenotypic profile of an exemplary aSNV in *SSH2*, we contrasted identified carriers with a curated set of non-carriers, homozygous for the derived allele, assessing a range of phenotypic traits. Given the especially dramatic effects of the *TKTL1* aSNV on neurogenesis reported by Pinson et al. in their cellular and animal models (*10*), we also included this high-frequency human-specific change in our investigations. Specifically, we identified carriers of the archaic *TKTL1* allele and used the available neuroimaging data (*17*) to study putative effects of the aSNV on brain morphology and cognitive traits. We use our findings to make recommendations about how to optimize future biobank-based investigations of human evolution.

### 103 carriers of archaic SNVs in 17 positions identified in UK Biobank across different ancestries

Based on the Kuhlwilm & Boeckx (*2*) catalogue of single nucleotide changes that distinguish modern humans and archaic hominins, we curated a list of 42 fixed missense changes with an allele frequency of one (AF = 1) at the time of publication, indicating complete fixation within the investigated modern human populations (see Methods, table S1). After quality control (see Methods) we then queried the whole-exome sequencing data of approximately 455,000 individuals (*18*–*20*) of the UKB to identify possible carriers of the archaic allele at 37 positions. We investigated four ancestry superclusters: European, African, East & South Asian (fig. S1A). In total, we identified 103 unique individuals carrying 118 aSNVs in 13 protein-coding genes (Table 1). All were heterozygous carriers, except for a female carrying a homozygous aSNV in *GRM6* (chr5:178994530), a gene encoding the ON bipolar metabotropic glutamate receptor, which overall also represents the genomic position with the largest carrier count.

**Table 1:**
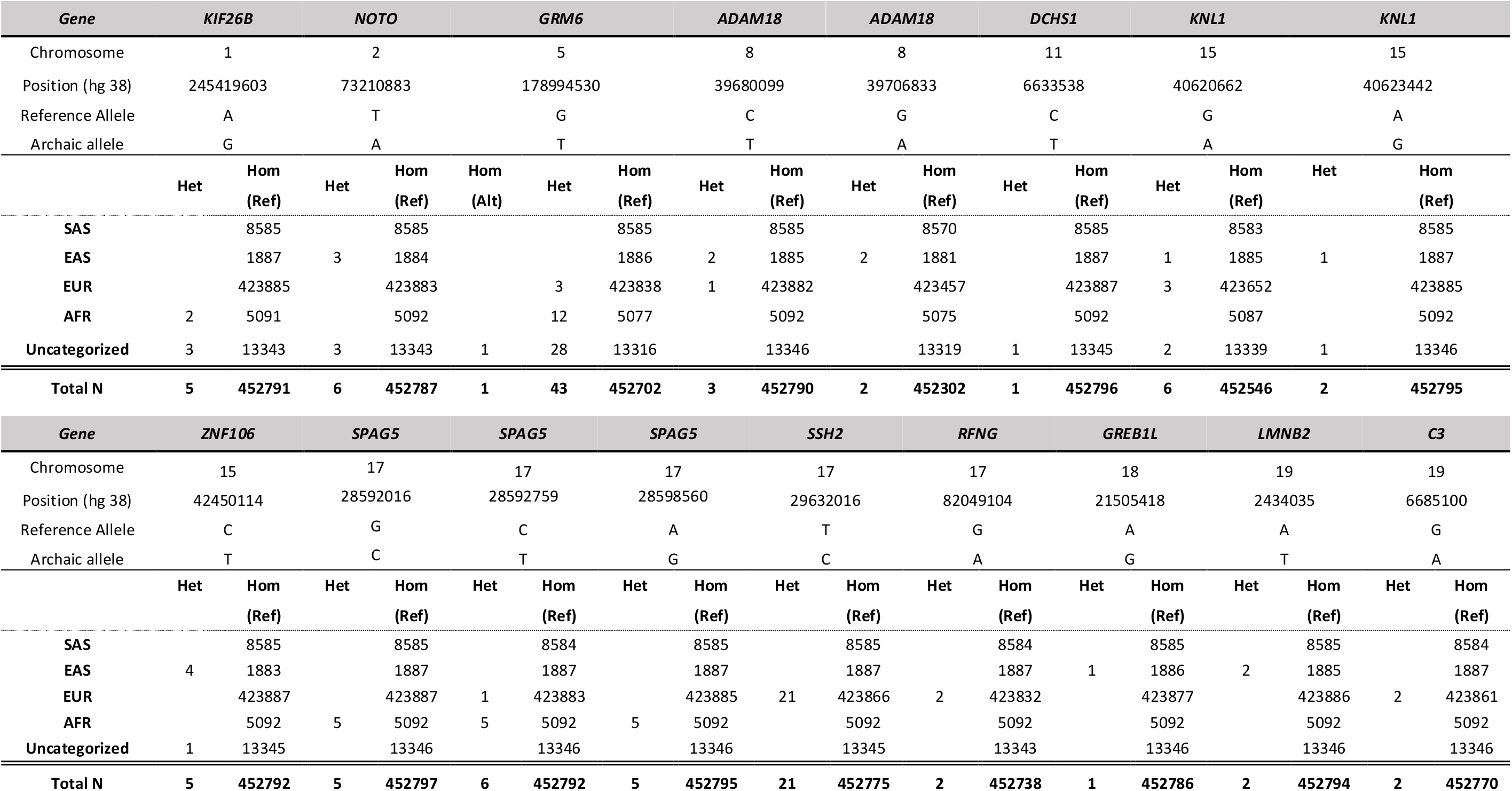
Overview of identified aSNV carriers in UK Biobank. Genotype count of carriers of each aSNV and respective individuals homozygous for the derived allele are noted per ancestry supercluster. Genomic positions are based on hg38; Het = Heterozygous, Hom = Homozygous, Ref = reference allele; SAS = South Asian; EAS = East Asian; EUR = European; AFR = African.

We observed diverging carrier counts for aSNVs across ancestry superclusters. Even though the UKB is comprised of predominantly European ancestry individuals (*16*) with only around 0.5% individuals of African and 1% of East Asian descent, nearly equal numbers of aSNV carriers were identified in the European, East Asian and African ancestry superclusters, highlighting allele frequency differences for these rare variants, and a bias towards European data being used previously to identify aSNVs.

We identified five individuals carrying a combination of three aSNVs in *SPAG5* (chr17:28,592,759; chr17:28,598,560; chr17:28,598,560), and two pairs of carriers who carry a combination of two aSNVs on *ADAM18* and *KNL1*, respectively (*ADAM18*: chr8:39,680,099; chr8:39,706,833; *KNL1*: chr15:40,620,662; chr15:40,623,442). In each case the aSNVs found in the same carriers were in tight linkage disequilibrium, thus were likely inherited together.

We also queried the relatedness status (up to 3^rd^ degree) of identified carriers and found only one related pair carrying an aSNV in *SSH2*. Thus, it is unlikely that the allele counts of identified aSNVs in our study are inflated due to relatedness.

### Phenotypes assessed in carriers of the *SSH2* aSNV do not deviate from matched non-carriers

Next, we showed how availability of biobank trait data can be used to query whether aSNVs have major phenotypic consequences in living humans. We chose an aSNV in *SSH2* (chr17:29632016) to exemplify this, since the encoded protein is a protein phosphatase with enzymatic properties regulating actin filament dynamics and possible functions in neurite outgrowth (*2, 21*–*23*), and because the variant was found in a relatively large number of unrelated carriers within a strict ancestry cluster (N = 19, see Methods). We chose the following traits for phenotypic assessment: body composition measures (body mass index, whole body fat mass), height, overall health rating, smoking status and highest qualification level as an indication of educational attainment. These phenotypes were selected a priori based on previously identified GWAS trait associations of *SSH2* (*24*) that further overlapped with traits linked to Neandertal admixture (*1, 25*–*29*).

For all continuous traits, carriers of the *SSH2* aSNV were within the standard trait distribution based on a matched set of individuals homozygous for the derived alleles (non-carriers; see Methods), and did not show a trend towards extreme values (Fig. 1A). A similar pattern was observed for categorical traits, where carriers show no strong deviating pattern from the matched non-carrier cohort (Fig. 1B-C). Given its putative roles in neurite outgrowth, prior associations of common variants with a broad array of brain imaging metrics (*24*), and the general involvement of protein phosphatases in psychiatric and neurological disorders (*30*), we tested the possible consequences of carrying aSNVs in *SSH2* for a range of neuropsychological traits. We did not observe diverging patterns for aSNV carriers compared to the non-carrier group (fig. S2).

**Figure 1:**
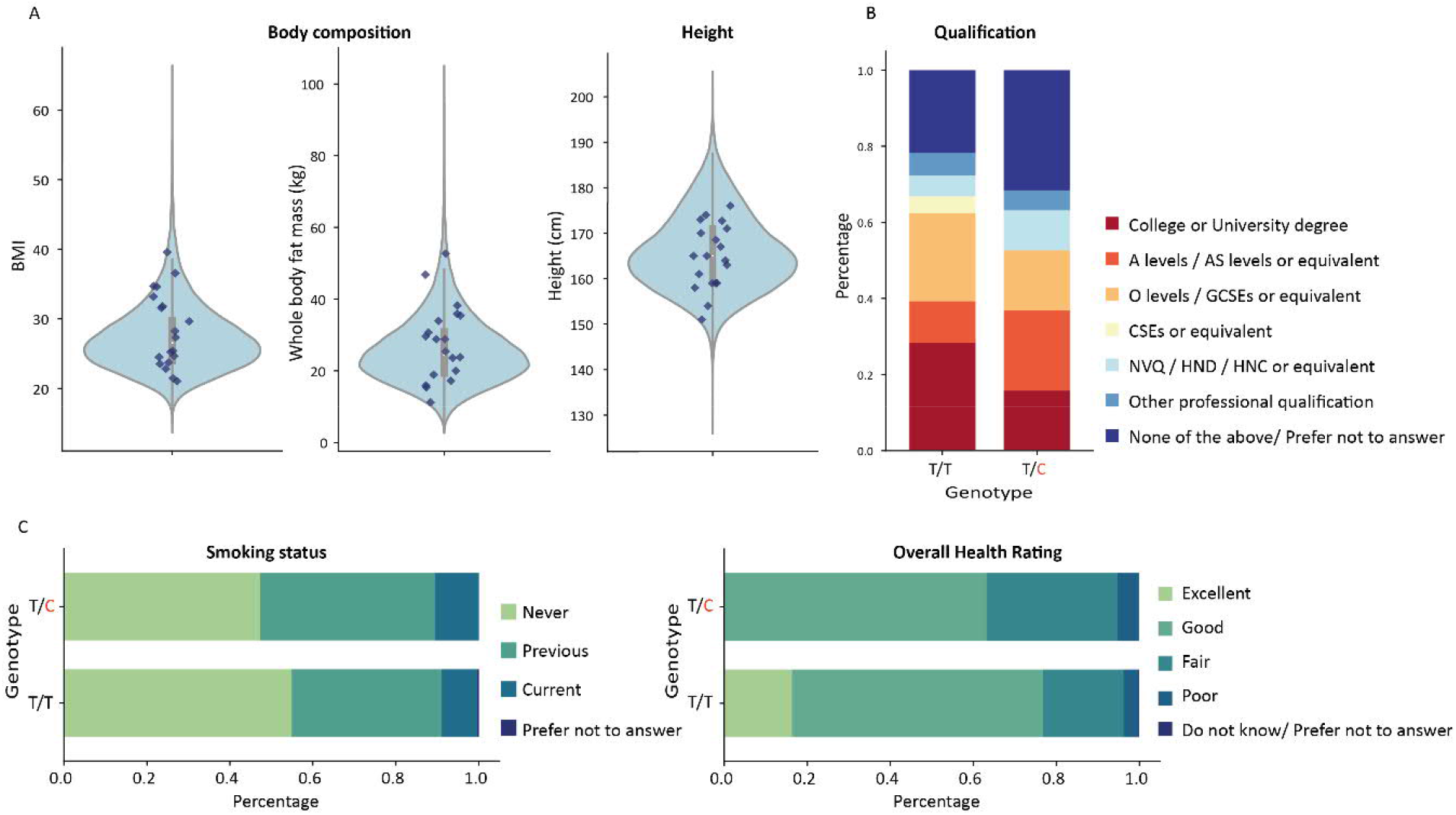
Investigating phenotypic effects in carriers of the *SSH2* aSNV. (A) Values of continuous traits are shown for each aSNV carrier as dark blue diamonds. Violin plots show the phenotypic distribution of matched set of non-carriers, with boxplots indicating the 25^th^ and 75^th^ percentiles, and whiskers representing 1.5 times the inter quartile range (IQR); (B & C) Stacked bar plots showing the percentage of highest qualification level, as well as health related measures for each genotype: T/T for matched non-carriers and T/C for aSNV carriers (N_aSNV_ = 19; N_Non-carrier_ = 39,501). A level = Advanced level, AS level = Advanced Subsidiary level, O level = Ordinary level, GCSE = General Certificate of Secondary Education, CSE = Certificate of Secondary Education, NVQ = National Vocational Qualification, HND = Higher National Diploma, HNC = Higher National Certificate.

### The archaic allele in *TKTL1* shows little consequence for frontal pole morphology and overall cognition in adult humans

We went on to study the archaic allele (A) of the rs111811311 polymorphism (A/G) of the *TKTL1* gene, located on the X chromosome. This missense change (yielding a lysine-to-arginine change at residue 317 of the long isoform) gained considerable prominence in recent literature when it was proposed by Pinson et al. as a major driver of human/Neandertal brain differences in evolution based on an array of functional experiments (*10*). Note that the variant was not among our curated list of aSNVs above, since it did not fit the criteria of full fixation in Kuhlwilm & Boeckx (*2*) (AF = 1), while a critique of Pinson et al. (*10*) has highlighted the existence of rs111811311 archaic allele carriers in gnomAD (*12*), but without any phenotypic follow-up. Querying the UKB resource for the *TKTL1* archaic allele, we identified 45 heterozygous and one homozygous female carrier, as well as 16 hemizygous male carriers across multiple ancestry groups (Table 2, fig S1C). Among these 62 carriers, we identified four pairs with a kinship coefficient below 0.042, indicating again that relatedness (up to the 3^rd^ degree) is unlikely to explain the larger number of carriers. One individual was identified with an archaic allele in both *TKTL1* and *KIF26B*.

**Table 2:**
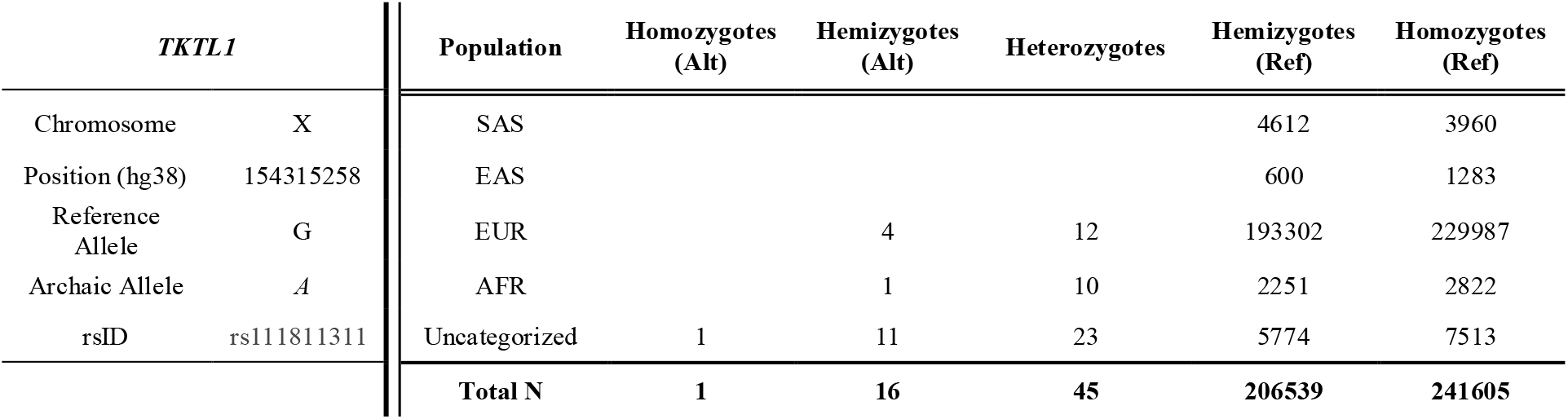
Overview of identified aSNV carriers for *TKTL1* in UK Biobank. Genotype count of carriers of the archaic allele and individuals homozygous or hemizygous for the derived allele are noted per ancestry supercluster. Genomic positions based on hg38; REF = reference allele; ALT = Alternative/Ancestral allele;

| *TKTL1* |  | Population | Homozygotes (Alt) | Hemizygotes (Alt) | Heterozygotes | Hemizygotes (Ref) | Homozygotes (Ref) |
| --- | --- | --- | --- | --- | --- | --- | --- |
| Chromosome | X | SAS |  |  |  | 4612 | 3960 |
| Position (hg38) | 154315258 | EAS |  |  |  | 600 | 1283 |
| Reference Allele | G | EUR |  | 4 | 12 | 193302 | 229987 |
| Archaic Allele | A | AFR |  | 1 | 10 | 2251 | 2822 |
| rsID | rs111811311 | Uncategorized | 1 | 11 | 23 | 5774 | 7513 |
|  |  | **Total N** | **1** | **16** | **45** | **206539** | **241605** |

Given that the cellular/animal work of Pinson et al. (*10*) linked the human-derived allele of *TKTL1* to substantial increases in neuron production in the prefrontal cortex, we contrasted imaging derived structural brain metrics of the frontal lobe in unrelated aSNV carriers (N = 5) and matched non-carriers, homozygous for the derived allele (N = 2145) to investigate the effects of carrying an archaic allele on frontal lobe surface area and cortical thickness in living human adults (Fig. 2).

**Figure 2:**
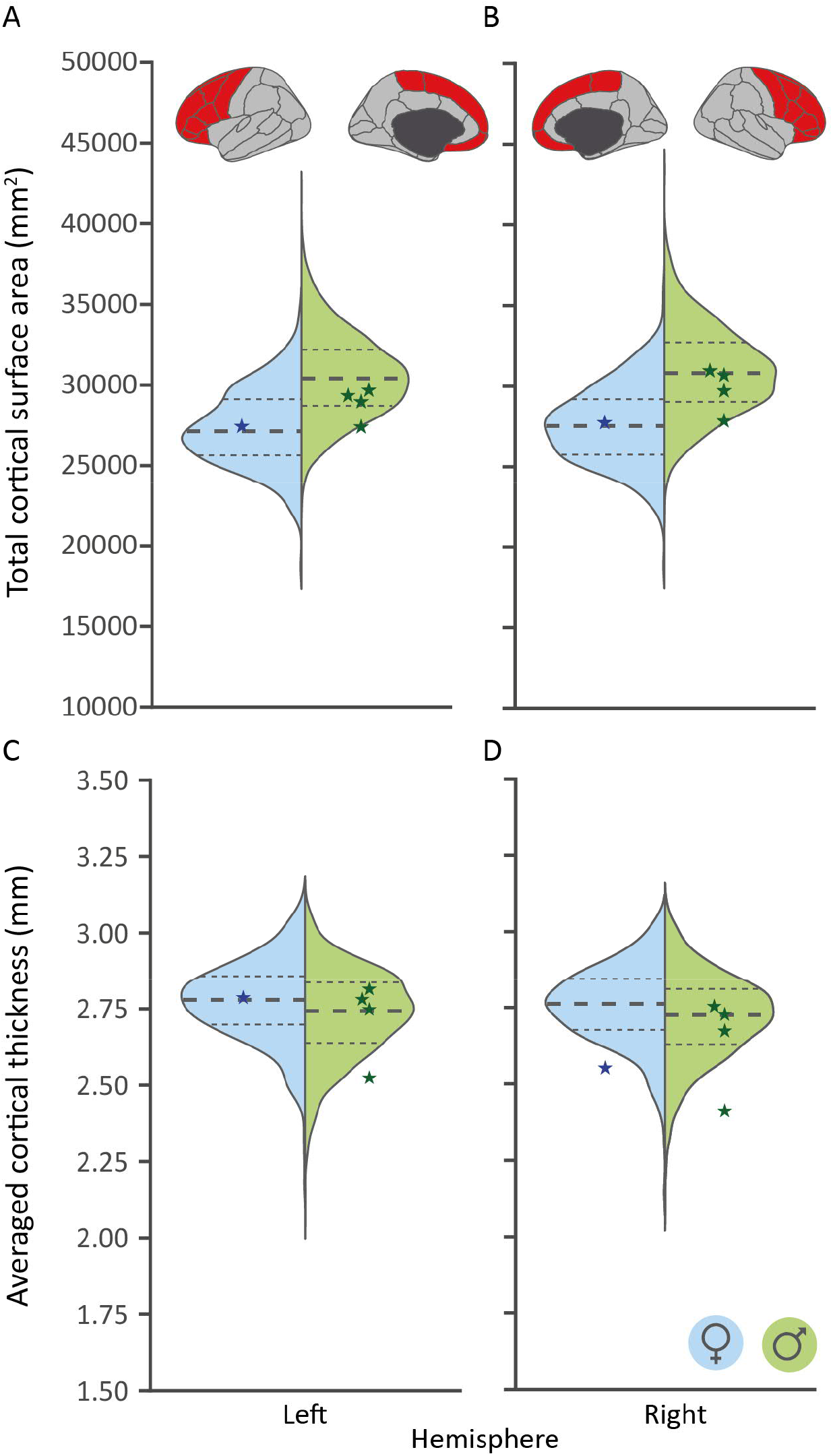
Carriers of the archaic allele of the *TKTL1* aSNV show no diverging cortical measures compared to a matched set of non-carriers. Archaic allele carrier values for each sex and metric are depicted as diamonds. These are overlaid over split violin plots indicating the phenotypic variability for both the female (left, blue) and male (right, green) matched non-carrier sample for total cortical surface area for both left and right hemisphere (A and B, respectively) and averaged cortical thickness (C and D, respectively); G: Reference/derived allele, A: Archaic allele. Tick dotted line indicates the median, while the thin dotted lines highlight both the 25^th^ and 75^th^ percentiles.

We found that the range of phenotypic variation of aSNV carriers lies in general within the 25^th^ and 75^th^ percentiles of the non-carriers for all cortical measures. This is in stark contrast with the pronounced effects shown in the various functional assays performed by Pinson et al. (*10*) which would predict substantial reductions in prefrontal cortex brain metrics of carriers of the archaic allele. As a sensitivity analysis, we repeated this approach in an ancestry matched cohort of only European carriers (N = 3) and a matched non-carrier cohort (n = 30), and obtained an even clearer overlap in phenotypic distributions (fig. S3).

Increased neuronal proliferation and expansion of the neocortex along the lineage leading to modern humans is argued by some to have been a driver of increased cognitive capacities of our species (*31, 32*). Indeed, based on the Pinson et al. (*10*) findings, some other researchers and commentators have proposed that the *TKTL1* protein-coding aSNV contributed to differences in cognition between *Homo sapiens* and extinct archaic humans (*33*). Thus, we also assessed educational qualification levels of carriers of the archaic *TKTL1* allele (N = 30) compared to matched non-carriers (N = 600) (Fig. 2). Due to the difference in zygosity, this was done separately for males and females.

**Figure 2:**
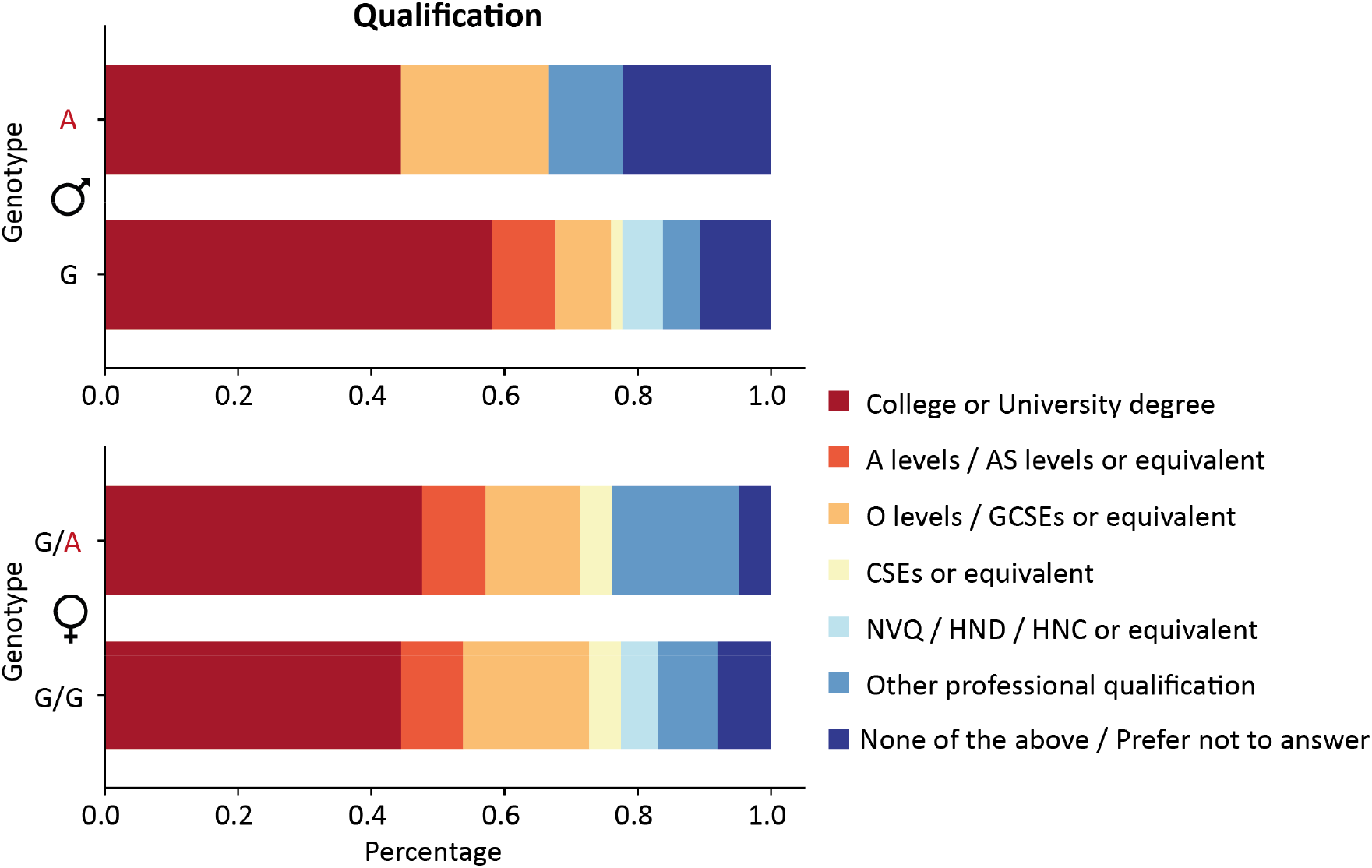
Qualification levels of carriers of archaic alleles of the *TKTL1* aSNV are similar to matched set of non-carriers. Stacked bar plots showing the percentage of highest qualification level by genotype. (Male sample: N_aSNV_ = 9; N_Non-carriersl_ = 180; Female sample: N_aSNV_ = 21; N_Non-carriers_ = 420); A level = Advanced level, AS level = Advanced Subsidiary level, O level = Ordinary level, GCSE = General Certificate of Secondary Education, CSE = Certificate of Secondary Education, NVQ = National Vocational Qualification, HND = Higher National Diploma, HNC = Higher National Certificate;

While the percentage of males with the highest qualification level was slightly lower for those with the archaic allele of *TKTL1*, it is striking that in both sexes a substantial proportion of carriers of this allele have a college or University degree. In particular, > 44% of males with only an archaic allele on this polymorphic site of the X chromosome have a college/University degree. This pattern of findings casts doubt on the idea that the human-derived change in *TKTL1* was a key player in the evolution of enhanced human cognitive abilities (*33*).

## DISCUSSION

This study brings a novel source of empirical data to questions regarding evolutionary impacts of protein-coding variants that distinguish between modern humans and our extinct archaic cousins, adding to the rich prior literature in this area, recently reviewed by Zeberg and colleagues (*3*). Our work identified 165 unique carriers of archaic SNVs for 18 out of a total of 38 interrogated genomic positions in around 450,000 individuals with exome data in UKB. Regarding phenotypic consequences of an exemplar aSNV in *SSH2*, one for which relatively large numbers of carriers were available, all interrogated traits fell within the typical range of variation, with no obvious divergence from the norm. A similar pattern was observable for *TKTL1* for frontal lobe structural measures as well as overall qualification level, despite this variant previously showing large effects on neocortical development in cellular/animal models.

Ever since the first high-coverage genome sequence of a Neandertal resulted in a catalogue of fixed missense aSNVs (*7*), the overall number has continually decreased, as more high quality Neandertal genomes and ever-increasing population databases of present-day humans have become available. For such protein-coding changes, the present study reduces the number of potential fully fixed genomic positions described in Kuhlwilm and Boeckx (*2*) that we investigated here from 37 to only 20, while the true number is likely even smaller. This raises questions over whether (some of) the aSNV carriers are explained by rare back-mutations, whether these sites were never fixed to begin with, or whether the ancestral allele was reintroduced post-fixation during admixture events (*3*). While for some genomic positions only a handful of carriers were found in the UKB, some positions present with higher carrier counts that make back-mutations an unlikely explanation (*1*). Further, higher ancestral allele counts were often evident in non-European ancestry groups. High genetic diversity within African populations (*34*) might partially explain this pattern, but considering the skewness of UKB towards White European ancestry (*16*) it remains intriguing. While it is known that some isolated populations have higher levels of archaic ancestry, either because they persisted since a common ancestor, as seen in the Khoe-San (*35*) or due to relatively recent admixture with Neandertals/Denisovans (e.g., Oceanian populations) (*36, 37*), there is no detailed catalogue of fixed human-specific changes across a range of ancestries that could be used as a reference point, given that most results of genomic studies are solely based on populations with European ancestry (*3*).

The presence of aSNV carriers in population databases, however, does not rule out the possibility that these DNA changes contributed to the formation of anatomically modern humans. While experimental validation in model systems is crucial to understand the impact of variation at these genomic positions, current approaches are laborious, with a range of known pitfalls (*12, 38*). The availability of phenotypic data in UKB makes it possible for the first time to query possible phenotypic consequences in present-day adult living humans that carry the variants of interest. As contrasting a range of traits of interest indicated no systematic differences between matched individuals homozygous for the derived allele and aSNV carriers, this could be seen as evidence that there are no major phenotypic consequences of carrying an ancestral rather than a derived version at the queried position. However, while the aSNV on *SSH2* was carefully chosen because of its potential effects on the enzymatic properties of the encoded protein, increasing the likelihood of observable phenotypic consequences, and based on the relatively large number of identified carriers, the possibility remains that we did not have a sufficient sample size to detect trait differences (*3*). Of note, all individuals were heterozygous carriers and still had one copy of the derived allele, therefore not reflecting the homozygous state observed in the Neandertal genome. Moreover, lacking power for a large phenome-wide screen, we chose phenotypes to target a priori based on broader literature, and might have thus missed a trait that is truly impacted by the variant. This raises a larger question of importance for the field: which phenotype(s) would best represent ‘the human condition’ in investigations of this kind? Latest archaeological evidence increasingly suggests cognitive and behavioural similarities with our extinct archaic cousins, meaning that differences, especially for complex traits, may well be subtle (*2, 4*–*6*). The lower carrier numbers of other aSNVs (at least within ancestry clusters) further limit the scope of currently feasible phenotypic investigations. A large phenome-wide scan sensitive enough to detect small deviations from the norm might highlight the most important phenotypes, as well as clarifying contributions of these genomic positions, but this will only be feasible when even larger sample sizes are available than at present.

While reported as only a human-specific high-frequency variant by Kuhlwilm and Boeckx (*2*), several reasons led us to include the aSNV in *TKTL1* in the current investigation. Firstly, the phenotypic consequences of ancestral versus derived alleles of this aSNV are well described in Pinson et al., based on their experiments in animal/cellular models (*10*), allowing for a more targeted phenotypic selection. Complementing findings from these models, the researchers also reported that disrupting *TKTL1* expression in fetal human brain neocortical tissue significantly reduced basal radial glial progenitors (*10*). Secondly, the effect sizes of the aSNV allele reported in Pinson et al. (*10*) were substantial, indicating that even with only a small number of identified carriers there should be good prospects of detecting such phenotypic consequences. Thirdly, the position of the aSNV on the X chromosome should lead to even more pronounced effects in males, who are hemizygous for either a derived or ancestral allele. Still, we saw no differences for carriers and matched sample of non-carriers in neuroanatomical properties of the frontal lobe even in male carriers, and a substantial proportion of these had a college/university degree, arguing against a major impact on cognitive functions. While the absence of consequences for adults might possibly be explained by compensatory mechanisms with mitigating effects on the developing frontal lobes, our results show that effect sizes identified in functional assays and model organisms cannot be directly extrapolated to the consequences of carrying these changes for adult human phenotypes (*15*).

Beyond general challenges related to rare variant analysis and the choice of target phenotypes, as discussed above, limitations of the current study include those related to the nature of the UKB cohort (restricted age range, lack of diversity in ancestral background, existence of participation bias (*16, 39*)), and the need for more and better-quality archaic hominin genomes to understand the genetic variation patterns in their populations.

The findings here resonate well with the recent perspective of the field set out by Zeberg and colleagues (*3*). With our concrete demonstration of biobank analyses, we provide new impetus towards promising avenues for future investigations: i) The inclusion of more large-scale diverse population databases (*40*–*43*) together with the information from the third high quality Neandertal genome (*9*) (and additional archaic genomes that might be sequenced) will likely yield a more representative catalogue of human-specific changes to help reconstruct how natural selection, archaic gene flow, and our demographic history together shaped our genome (*1, 3, 34, 44*). ii) Given the ever-decreasing number of such sites, it seems warranted to abandon the notion of fully-fixed variants, broadening the scope to also take high-frequency non-fixed changes into account. Kuhlwilm and Boeckx (*2*) already made a start in this direction and expanded their catalogue accordingly, but with the availability of larger and more diverse databases, this list will need updating. As more population databases are also including whole genome sequencing, the search can be expanded further to include high-frequency changes in regulatory regions (*45*). iii) Our results indicate that looking at each of these genomic positions individually might not be so informative and that future work focusing on their aggregated effects could be valuable (*2, 3*). One way to achieve this would be by grouping high-frequency changes according to their potential functions [see Kuhlwilm and Boeckx (*2*) for an initial categorization]. Further, a list of high-frequency variants could also be used for burden testing, which would additionally allow formal statistical analyses of possible effects (*46, 47*).

Overall, by leveraging the availability of archaic variation in modern biobanks, our study has provided evidence against the notion of fixed genomic changes on the human lineage, highlighted that individual interrogation of the key sites is unlikely to yield major insights into the emergence of complex human traits, and emphasises again the importance of including diverse ancestral backgrounds in studies on the origins of our species.

## Acknowledgments

This research was conducted using the UK Biobank resource under application no. 16066 with CF as the principal applicant. Our study made use of brain imaging derived phenotypes and pre-processed imaging data generated by an image processing pipeline developed and run on behalf of UK Biobank. SEF is a member of the Center for Academic Research and Training in Anthropogeny.

## Funding

Max Planck Society core funding (BM, MLA, GA, EE, DS, CF, SEF)

Dutch Research Council NWO; VI.Veni.202.072 (EE)

## Author contributions

Conceptualization: BM, MLA, EE, GA, SEF

Resources: CF, SEF

Methodology: BM, EE, GA, DS, SEF

Data analysis: BM, MLA

Writing – original draft: BM

Writing – review & editing: BM, MLA, EE, GA, DS, CF, SEF

## Competing interests

Authors declare that they have no competing interests.

## Data and materials availability

Whole exome sequencing data, imaging-derived phenotypes and other phenotypic data used in this study are available from UK Biobank (https://www.ukbiobank.ac.uk). All scripts used for the analyses are available on the project GitLab repository (https://gitlab.gwdg.de/barmol/fixedVariant).

## Supplementary Materials

Materials and Methods

Figs. S1 to S3

Tables S1

References (*48*–*53*)

## Notes

### Competing Interest Statement

The authors have declared no competing interest.

### Summary of Updates

Integrated pre-submission feedback throughout the manuscript

