## Supplementary Material for "Evaluating the effects of archaic protein-altering variants in living human adults"

**The PDF file includes:**

Materials and Methods
Figs. S1 to S3
Table S1

### Materials and Methods

Dataset

All data used were obtained from the UK Biobank (UKB) under the research application 16066 with Clyde Francks as the principal investigator. Detailed descriptions of the data used as well as sample, genotype and variant specific quality control (QC) are given below. The UK Biobank has received ethical approval from the National Research Ethics Service Committee North West-Haydock (reference 11/NW/0382) and all of their procedures were performed in accordance with the World Medical Association. Informed consent was obtained for all participants by UKB with details about data collection and ethical procedures described elsewhere (*16*, *48*).

Whole exome sequencing data

Whole-exome sequencing was performed and data were processed by the UKB according to protocols described elsewhere (*18–20*). Briefly, the samples were multiplexed and then sequenced using 75-base-pair paired-end reads with two 10-base-pair index reads on the Illumina NovaSeq 6000 platform using either S2 (first exome release) or S4 flow cells. Sample-specific FASTQ files, representing all the reads generated for that sample, were mapped to the GRCh38 genome reference using the BWA-meme algorithm 60. Subsequently, the binary alignment files (BAM) for each sample contained the mapped reads' genomic coordinates, quality information, and the degree to which a particular read differed from the reference at its mapped location. Duplicated reads were removed with the Picard (*49*) MarkDuplicates tool. GVCF files were then produced using the weCall variant caller. Upon completion of variant calling, individual sample BAM files were converted to fully lossless CRAM files using samtools (*50*).

For this project, we made use of the Broad 455k exome gnomAD VCF files (UKB data field 24068): population VCF files that have been returned to UK Biobank as part of the 'alternative exome processing' (UKB Category 172). Here, original UKB CRAM files were re-processed according to the GATK Best Practices, aligning reads using BWA-MEM 0.7.15.r1140 and processing reads using Picard and GATK with protocols described in detail in Karczewski et al.(*51*)

Variant selection

We based our selection of fixed, amino acid changing SNVs on a list of high-frequency human-specific missense changes described in Kuhlwilm & Boeckx (*2*), where only those SNVs with an allele frequency of one were selected, indicating total fixation at the time of their publication. These 42 genomic positions were lifted to GRCh38 using Liftover (https://liftover.broadinstitute.org/). We further confirmed that each SNV was indeed located in a translated exon using gnomAD v3.1.2. (https://gnomad.broadinstitute.org/) and the UCSC Genome Browser (https://genome.ucsc.edu/), which lead to exclusion of 3 variants located in *C1orf159* (chr1:1091245), *DNHD1* (chr11: 6534188) and *DNMT3L (*21: 44251169). One genomic position in *TBC1D3* (chr17:38202786) was excluded due to ambiguous results after liftover to GRCh38.

Due to its reported profound effects on brain development in cellular/animal models (*10*), a SNV in *TKTL1* on chromosome X (X: 154315258), previously reported as fixed by Prüfer et al. (*7*), was also included in our analysis despite being reported as a high-frequency variant (AF: 0.999694) in Kuhlwilm & Boeckx (*2*).

This resulted in a total of 39 SNVs in 33 genes that were put forward for further analysis (see table S1).

Sample specific quality control

On all available individuals included in the gnomAD VCFs (N = 454,672), we first applied sample level quality control measures. This entailed excluding individuals with a mismatch of their self-reported (UKB data field 31) and genetically inferred sex (UKB data field 22001), as well as individuals with putative aneuploidies (UKB data field 22019), or individuals who were determined as outliers based on heterozygosity (PC corrected heterozygosity >0.1903) or genotype missingness rate (missing rate >0.05) (UKB data field 22027), leading to a final sample of 452,797 individuals.

Variant and genotype QC

For further analysis we moved to the UK Biobank Research Analysis Platform (UKB RAP, https://ukbiobank.dnanexus.com) and queried the curated list of genomic position detailed above using bcftools (version 1.17) (*52*) to identify possible carriers of the archaic allele.
To assure that identified carriers did not represent sequencing errors, we only included archaic SNVs who were called with PASS in the VCF. This variant filter is based on a combination of a random forest classifier and hard filters, detailed in Karczewski et al. (*51*). Only one SNV (chr9:6606647) did not pass these filters and was discarded for further analysis. We further used Hail (https://github.com/hail-is/hail) as implemented in JupyterLab on the UKB RAP for quality control of individual genotype data, where genotypes at the specific positions were filtered based on genotype quality (QUAL> 20), depth (DP> 10), and allele balance for heterozygous genotypes (AB > 25% < 75%), leading to different sample counts per queried position.

Variant distribution per ancestry cluster

We inferred ancestry for all individuals surviving the quality control outlined above. We first used the self-reported ethnicity (*16*) (UKB data field 21000-0.0) as provided by UKB and grouped each individual into 4 major ancestry clusters [European (EUR), African (AFR), South Asian (SAS), East Asian (EAS)]. Individuals who reported ‘Mixed’, ‘Other’, ‘Do not know’ and ‘Do not want to answer’ were grouped as ‘Uncategorized’. We then used the first four provided genetic ancestry principal components (UKB data field Data-Field 22009-0.1 - 22009-0.4) and assigned each individual to one of the respective major ancestry clusters using hard cut-offs. Individuals labelled as ‘Uncategorized’, which only showed a highly dispersed cluster, were reassigned to one of four superclusters if their PC’s fell within the respective boundaries. Otherwise, the individuals kept their initial category. This resulted in 5092 individuals within the AFR ancestry cluster, 1887 individuals within the EAS cluster, 423,887 individuals within the EUR ancestry cluster and 8585 individuals within the SAS cluster, while 13,346 individuals could not be assigned to one of the superclusters and were thus labelled as ‘uncategorized’.

Relatedness

To infer if relatedness could explain an accumulation of archaic SNVs at certain positions, we identified if carriers had a kinship coefficient > 0.0442 (UKB data field 22021), but initially did not exclude any individuals based on relatedness. For phenotypic analysis, this information was used, and one individual from each pair of relatives was excluded, where we prioritised the exclusion of non-carriers, as well as individuals related to a larger number of other individuals.

**Phenotypic analyses**

<u>*SSH2*</u>

For phenotypic analysis, we chose one exemplary genomic position for a qualitative comparison of carriers to non-carriers. Our initial query of the exome data highlighted a group of 21 individuals with an aSNV on chr17:29632016 within *SSH2,* where all 21 were grouped within the EUR ancestry cluster, and 20 were also assigned to the more stringent 'White British' ancestry (UKB data field 22006) (*16*). We further excluded one carrier due to relatedness, leading to a final carrier count of 19 unrelated, White British individuals (14 female; mean age ± SD: 59.42 ± 7.94 years).

We derived an age- (UKB data field 21003-0.0), sex- (UKB data field 31-0.0) and ancestry-matched (white British, UKB data field 22006) unrelated sample of individuals homozygous for the derived allele, where each carrier was paired with 2079 unique non-carriers. This led to a matched non-carrier cohort of 39,501 individuals (29,106 females, mean age ± SD: 60.69 ± 7.73 years).

<u>Trait selection</u>

Based on literature detailing the phenotypic legacy of previous admixture events, and previous genetic associations with *SSH2* variants as detailed in the GWAS catalogue, we selected a range of traits to investigate potential phenotypic effects of carrying an aSNV: BMI (UKB data field 21001-0.0), Whole-body-fat-mass (UKB data field 23100-0.0), Height (UKB data filed 50-0.0), Overall Health rating (2178-0.0), and Qualification (UKB data field 6138-0.0). For each individual we included only the highest qualification level reported in the analysis. As the GWAS catalogue highlighted several associations of *SSH2* with different brain phenotypes and given prior proposed links of archaic admixture to psychiatric disorders, we also included a broad range of neuropsychiatric metrics: "Seen doctor (GP) for nerves, anxiety, tension or depression" (UKB data field 2190-0.0), "Seen Psychiatrist for nerves, anxiety, tension or depression" (UKB data field 2100-0.0), Moodiness (UKB data field 1920-0.0), Miserableness (UKB data field 1930-0.0), loneliness (UKB data field 2020-0.0) and risk-taking (UKB data field 2040-0.0).

<u>*TKTL1*</u>

Given the profound effects of the derived allele of *TKTL1* on brain development in experiments described by Pinson at al. (*10*), we also included targeted investigations of genotype/phenotype relationships for this aSNV. As that prior study had highlighted increase neuronal counts specifically in the frontal lobe (*10*) we focused our phenotypic analysis first on relevant neuroanatomical data, making use of imaging-derived phenotypes generated by an imaging-processing pipeline developed and run on behalf of the UK Biobank (*17*, *53*). Imaging-derived structural measures (UK Biobank category 192) were available for 5 unrelated carriers (1 female; mean age ± SD: 71.2 ± SD 6.14 years). To estimate total frontal lobe surface area, we summed for each brain hemisphere the following imaging-derived phenotypes Superior Frontal (UKB data field 26748-2.0 & 26849-2.0), Rostral Middle Frontal (UKB data field 26747-2.0 & 26848-2.0), Caudal Middle Frontal (UKB data field 26724-2.0 & 26825-2.0), Pars Opercularis (UKB data field 26738-2.0 & 26839-2.0), Pars Triangularis (UKB data field 26740-2.0 & 26841-2.0), Pars Orbitalis (UKB data field 26739-2.0 & 26840), Lateral Orbifrontal (UKB data field 26732-2.0 26833-2.0) , Medial Orbifrontal (UKB data field 26734-2.0 & 26835-2.0), Precentral (UKB data field 26744-2.0 & 26845-2.0), Paracentral (UKB data field 26737-2.0 & 26838-2.0) and frontal pole (UKB data field 26752-2.0 & 26853-2.0). For the same cortical parcellations we used the imaging-derived cortical thickness (UKB data field 26782-2.0 & 26883-2.0, UKB data field 26781-2.0 & 26882-2.0, UKB data field 26758-2.0 & 26859-2.0, UKB data field 26772-2.0 & 26873-2.0, UKB data field 76774-2.0 & 26875-2.0, UKB data field 26773-2.0 & 26874-2.0, UKB data field 26766-2.0 & 26867-2.0, UKB data field 26768-2.0 & 26869-2.0, UKB data field 26778-2.0 & 26879-2.0, UKB data field 26771-2.0 & 26872-2.0, UKB data field 26786-2.0 & 26887-2.0) to estimate the averaged cortical thickness for each hemisphere in the frontal pole. As Pinson et al. (*10*) and others (*33*) have suggested that the change from archaic to human *TKTL1* could have played an important role for the evolution of complex behaviour, we also looked at qualification level (UK Biobank field 6138-0.0). For each individual we included only the highest qualification level reported in the analysis.

As *TKTL1* carriers were found in more than one ancestry cluster, the matched samples of non-carriers were set up as following.

- For brain imaging phenotypes, we first used all 5 carriers with imaging data and identified an equal number of individuals homozygous for the derived allele (N = 429) per carrier which were only matched by age (UK Biobank field 21003-2.0) and sex (UKB data field 31-0.0). This resulted in a non-carrier sample across ancestries of 2145 individuals (429 female; mean age ± SD: 71.2 ± SD 5.49 years).
- For a sensitivity analysis of the above, we only used the 3 European carriers (1 female, mean age ± SD: 73.67 ± SD 3.21 years), where 10 non-carrier individuals were matched to each carrier by the first two genetic principal components (PC1 & PC2 ± 2.5, respectively; UKB data field 22009-0.1 and 22009-0.2), age (± 2.5 years; UK Biobank field 21003-2.0) and sex (UKB data field 31-0.0). This resulted in 30 unique non-carriers (10 females; mean age ± SD: 73.2± SD 3.21 years).
- For our analysis of qualification level, we only selected unrelated carriers, where we could identify 20 unique, matched (PC1 & PC2 ± 2.5, respectively; age ± 2.5 years, sex) non-carriers, which led to a final sample comprised of 30 carriers (21 female; 11 AFR, 10 EUR, 9 UNC; mean age ± SD: 54.4 ± SD 7.67 years) and 600 matched non-carriers (420 females; mean age ± SD: 54.49 ± SD 7.73 years);

For all included qualitative traits used in this study we combined possible answer options ‘Do not know’, ‘Do not want to answer’ and/or ‘None of the above’ to one item.

**Fig. S1**

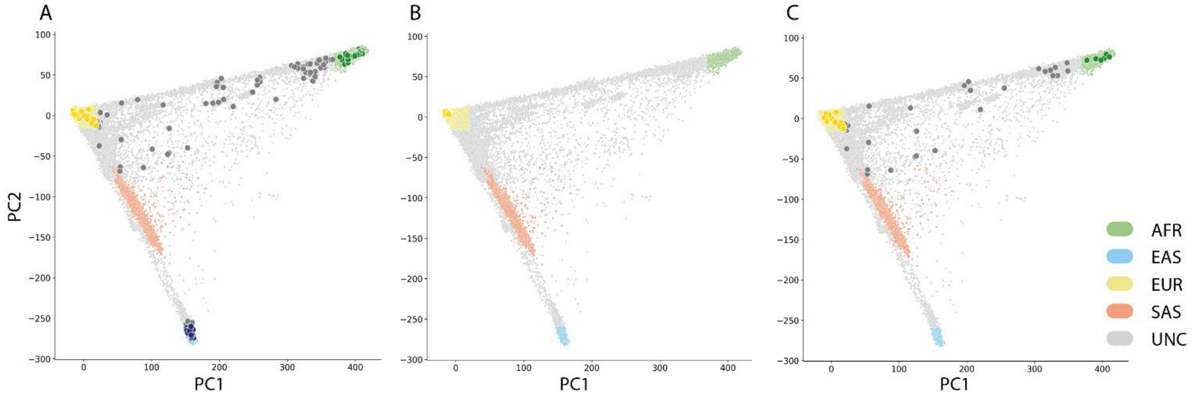


**Fig. S1. Ancestry distribution of identified aSNV carriers**. Top two principal components plotted for (A) all aSNV carriers (N = 165), (B) carriers of an aSNV in *SSH2* (N = 21), and (C) carriers of an aSNV in *TKTL1* (N = 62). Carriers are overlaid as darker shaded dots over individuals that do not carry an aSNV. Ancestry superclusters are colour coded and were defined based on the top 4 principal components using self-report ethnic background (UKB data field 21000); AFR = African, EAS = East Asian, EUR = European, SAS = South Asian, UNC = Unclassified.

### Fig. S2

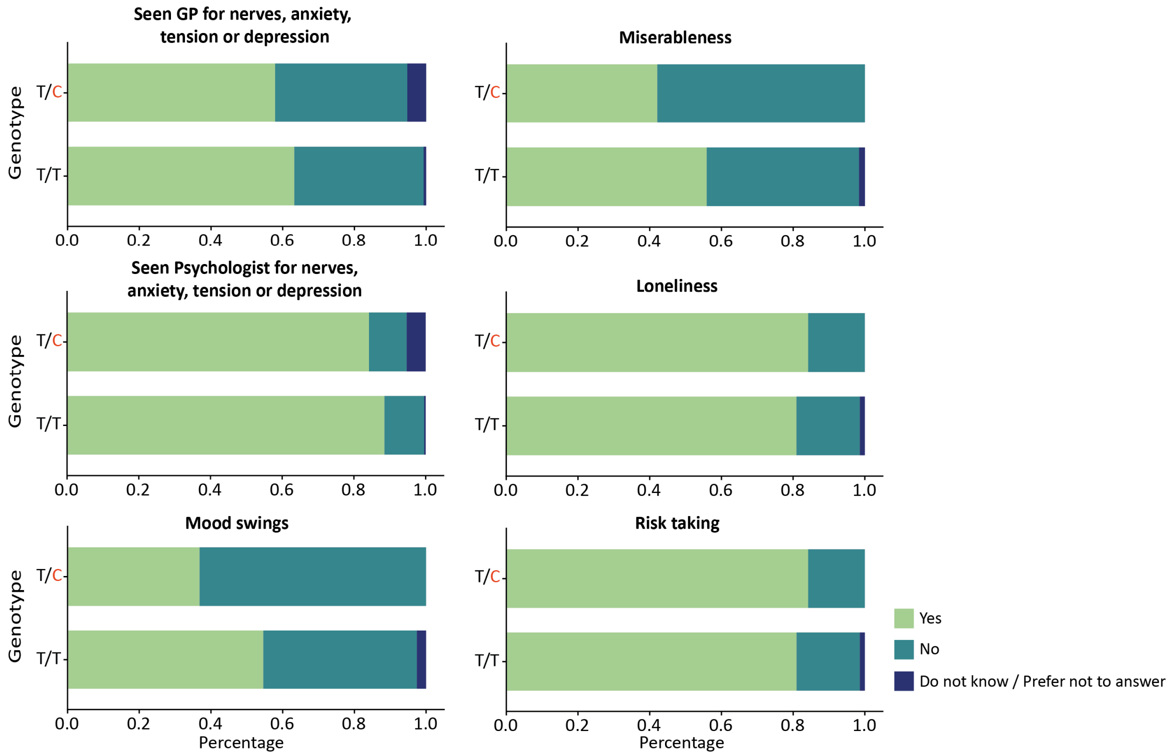


**Fig. S2. Neuropsychiatric traits in carriers of an archaic SNV in *SSH2*.** Stacked bar plots highlight the percentage of answers (Yes, No, do not know/Prefer not to answer) with respect to a set of categorical traits within the neuropsychiatry domain per genotype ($N_{aSNV}$ = 19; $N_{Non\text{-}carrier}$ = 39,501).

**Fig. S3**

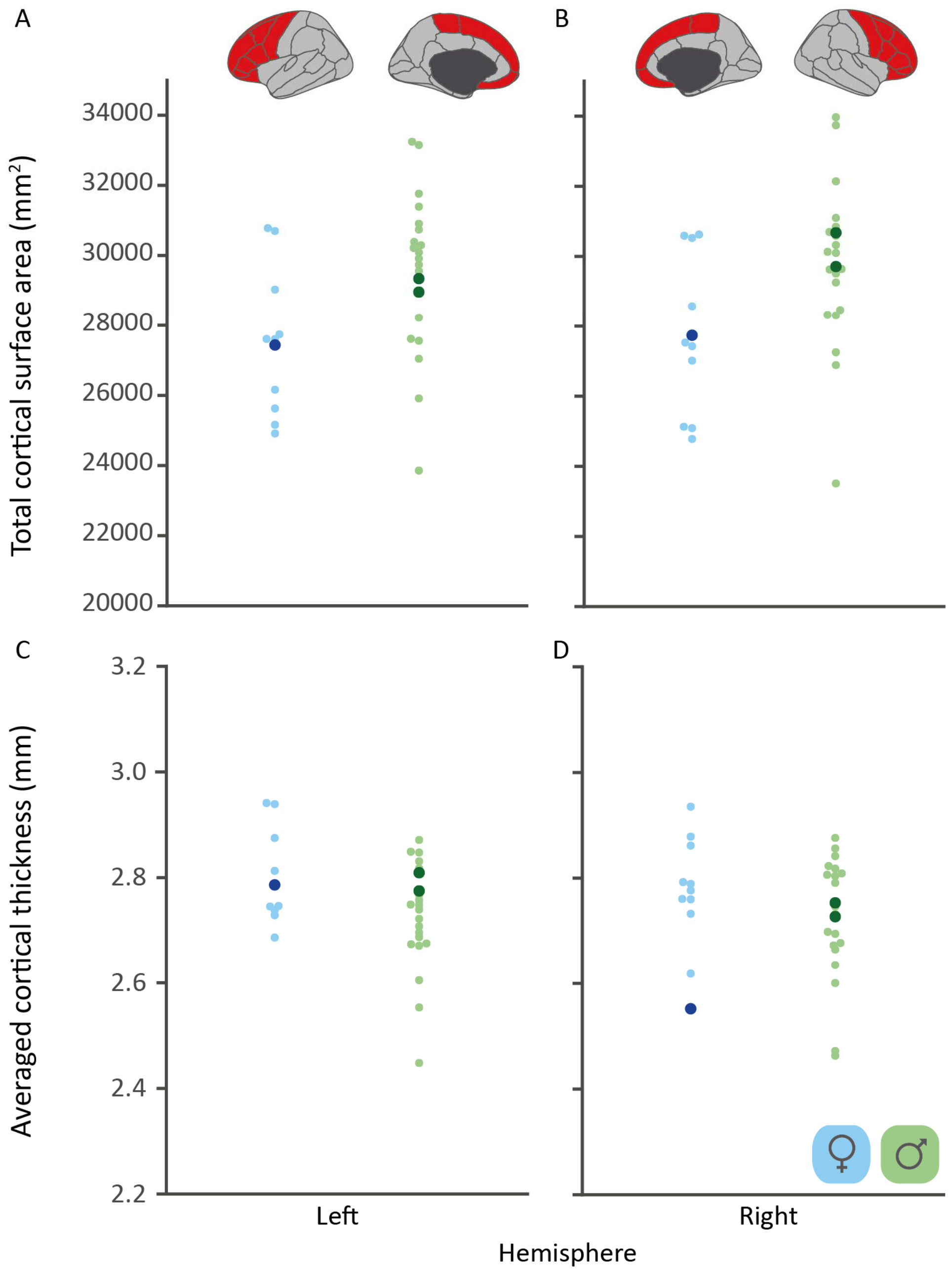


**Fig. S3**. **Carriers of the archaic allele of the *TKTL1* aSNV show no diverging cortical measures compared a matched set of non-carriers.** Strip plots depict the phenotypic variability in both female (blue) and male (green) matched samples of non-carriers for total frontal lobe cortical surface area for both left and right hemisphere (A and B, respectively) and averaged cortical thickness (C and D, respectively). Carrier values for each sex and metric are overlaid as darker shaded dots. In this sensitivity analysis, only participants with European ancestry were included ($N_{aSNV} = 3$, $N_{Non\text{-}carrier} = 30$)

### Table S1

| Gene | *C1orf159** | *GBP5* | *KIF26B* | *NOTO* | *PROM2* | *ANKMY1* |
|---|---|---|---|---|---|---|
| Chromosome | 1 | 1 | 1 | 2 | 2 | 2 |
| Position (Hg38) | 1091245 | 89262377 | 245419603 | 73210883 | 95279944 | 240524049 |
| rsID | rs868110071 | rs2100717044 | rs1284226189 | rs372916128 | rs773160830 | rs2152357551 |
| Reference allele | G | T | A | T | C | T |
| Archaic allele | A | G | G | A | G | G |
| Amino acid change | | p.Glu497Ala | p.Gly342Ser | p.Asn237Ile | p.Glu458Asp | p.Asn467Lys |
| Ensembl transcript | | ENST00000370459.8 | ENST00000407071.7 | ENST00000398468.4 | ENST00000317620.14 | ENST00000272972.7 |

| Gene | *SCAP* | *BOD1L1* | *GRM6* | *AHR* | *ADAM18* | *ADAM18* |
|---|---|---|---|---|---|---|
| Chromosome | 3 | 4 | 5 | 7 | 8 | 8 |
| Position (Hg38) | 47427659 | 13595914 | 178994530 | 17335768 | 39680099 | 39706833 |
| rsID | rs1706196218 | rs2108927976 | rs971140136 | rs1782348480 | rs931223234 | rs758696627 |
| Reference allele | A | C | G | T | C | G |
| Archaic allele | G | G | T | C | T | A |
| Amino acid change | p.Thr140Ile | p.Arg2684Gly | p.Thr139Pro | p.Ala381Val | p.Val565Ala | p.Lys649Arg |
| Ensembl transcript | ENST00000265565.10 | ENST00000040738.10 | ENST00000231188.9 | ENST00000242057.9 | ENST00000265707.10 | ENST00000265707.10 |

| Gene | *GLDC*** | *ANKRD30A* | *BBIP1* | *DNHD1** | *DCHS1* | *ZNHIT2* |
|---|---|---|---|---|---|---|
| Chromosome | 9 | 10 | 10 | 11 | 11 | 11 |
| Position (Hg38) | 6606647 | 37219713 | 110900521 | 6534188 | 6633538 | 65117241 |
| rsID | rs1818739987 | rs1368438340 | rs2134016025 | rs1852891266 | rs752702880 | rs2137212312 |
| Reference allele | G | A | G | T | C | T |
| Archaic allele | A | G | A | C | T | C |
| Amino acid change | p.Phe220Leu | p.Arg1334Gln | p.Ile92Thr | | p.Asn777Asp | p.Arg138Gln |
| Ensembl transcript | ENST00000321612.8 | ENST00000361713.2 | ENST00000454061.5 | | ENST00000299441.5 | ENST00000310597.6 |

| Gene | *ZNHIT2* | *FRMD8* | *PRDM10* | *KNL1* | *KNL1* | *ZNF106* |
|---|---|---|---|---|---|---|
| Chromosome | 11 | 11 | 11 | 15 | 15 | 15 |
| Position (Hg38) | 65117485 | 65387131 | 129902398 | 40620662 | 40623442 | 42450114 |
| rsID | rs2137212866 | rs2137854779 | rs1831882552 | rs755472529 | rs1033262852 | rs368058761 |
| Reference allele | A | A | T | G | A | C |
| Archaic allele | G | G | G | A | G | T |
| Amino acid change | p.Arg57Cys | p.Arg32His | p.Thr1133Asn | p.His133Arg | p.Gly1060Ser | p.Thr720Ala |

| | | | | | | |
|---|---|---|---|---|---|---|
| Ensembl transcript | ENST00000310597.6 | ENST00000531296.1 | ENST00000358825.9 | ENST00000399668.7 | ENST00000399668.7 | ENST00000564754.7 |

| Gene | *CDH16* | *SPAG5* | *SPAG5* | *SPAG5* | *SSH2* | *SSH2* |
|---|---|---|---|---|---|---|
| Chromosome | 16 | 17 | 17 | 17 | 17 | 17 |
| Position (Hg38) | 66913161 | 28592016 | 28592759 | 28598560 | 29632016 | 29632240 |
| rsID | rs2145458747 | rs1597598245 | rs768937910 | rs756182312 | rs761423117 | rs2150961500 |
| Reference allele | T | G | C | A | T | C |
| Archaic allele | C | C | T | G | C | T |
| Amino acid change | p.Ala342Thr | p.Asps410His | p.Glu162Gly | p.Pro43Ser | p.Gly1060Ser | p.Lys985Arg |
| Ensembl transcript | ENST00000299752.9 | ENST00000321765.10 | ENST00000321765.10 | ENST00000321765.10 | ENST00000540801.6 | ENST00000540801.6 |

| Gene | *TBC1D3**** | *RFNG* | *GREB1L* | *LMNB2* | *C3* | *SIX5* |
|---|---|---|---|---|---|---|
| Chromosome | 17 | 17 | 18 | 19 | 19 | 19 |
| Position (Hg38) | 38202786 | 82049104 | 21505418 | 2434035 | 6685100 | 45762030 |
| rsID | | rs1243956307 | rs976443505 | rs1441130454 | rs2145396648 | rs576741494 |
| Reference allele | T | G | A | A | G | T |
| Archaic allele | A | A | G | T | A | C |
| Amino acid change | | p.Tyr281His | p.Ser1360Asn | p.Met425Leu | p.Val1286Ala | p.Pro509Leu |
| Ensembl transcript | | ENST00000310496.9 | ENST00000424526.7 | ENST00000325327.4 | ENST00000245907.11 | ENST00000457052.3 |

| Gene | *NCOA6* | *DNMT3L** | *ADSL* | *OTUD5* | *SHROOM4* | *ZNF185* |
|---|---|---|---|---|---|---|
| Chromosome | 20 | 21 | 22 | X | X | X |
| Position (Hg38) | 34749726 | 44251169 | 40364974 | 48957495 | 50634436 | 152959706 |
| rsID | rs1474484023 | rs1412603589 | rs372051078 | rs2064271773 | rs1931232889 | rs1556908367 |
| Reference allele | A | T | T | T | G | A |
| Archaic allele | C | A | C | G | A | G |
| Amino acid change | p.Met823Ile | | p.Ala429Val | p.Leu26Met | p.Val546Ala | p.Ala473Thr |
| Ensembl transcript | ENST00000359003.7 | | ENST00000623063.3 | ENST00000156084.8 | ENST00000289292.11 | ENST00000449285.6 |

| Gene | *TKTL1* |
|---|---|
| Chromosome | X |
| Position (Hg38) | 154315258 |
| rsID | rs111811311 |
| Reference allele | G |
| Archaic allele | A |
| Amino acid change | p.Lys317Arg |
| Ensembl transcript | ENST00000369915.8 |

**Table S1. Overview of all genomic positions of interest**. Based on the Kuhlwilm & Boeckx[2] catalogue of single nucleotide changes that distinguish modern humans and archaic hominins, we include all putative fixed genomic locations with an allele frequency of one (AF = 1) at the time of publication, as well as the high-frequency change on *TKTL1*. Please note, rsIDs refer to the change from derived to archaic allele while amino acid changes indicate the the amino acid change that occurred after *Homo sapiens* split from its common ancestor (archaic_position_derived); *excluded position as main transcript was in an intron; **excluded as position did not pass variant quality control; ***excluded as position/change could not be unambiguously identified.
